# Redefining the spliceosomal introns of the sexually transmitted parasite *Trichomonas vaginalis* and its close relatives in columbid birds

**DOI:** 10.1101/2024.11.13.623467

**Authors:** Francisco Callejas-Hernández, Mari Shiratori, Steven A. Sullivan, Frances Blow, Jane M. Carlton

## Abstract

*Trichomonas vaginalis* infects the urogenital tract of men and women and causes the sexually transmitted infection trichomoniasis. Since the publication of its draft genome in 2007, the genome has drawn attention for several reasons, including its unusually large size, massive expansion of gene families, and high repeat content. The fragmented nature of the draft assembly made it challenging to obtain accurate metrics of features, such as spliceosomal introns. The number of introns identified varied over the years, ranging from 41 when first characterized in 2005, to 32 in 2018 when the repertoire was revised. In both cases, the results suggested that more introns could be present in the genome. In this study, we exploited our new *T. vaginalis* G3 chromosome-scale assembly and annotation and high-coverage transcriptome datasets to provide a definitive analysis of the complete repertoire of spliceosomal introns in the species. We developed a custom pipeline that distinguishes true splicing events from chimeric alignments by utilizing the extended motifs required by the splicing machinery and experimentally verified the results using transcript evidence. We identified a total of 63 active introns and 34 putative “inactive” intron sequences in *T. vaginalis*, enabling an analysis of their length distribution, extended consensus motifs, intron phase distribution (including an unexpected expansion of UTR introns), and functional annotation. Notably, we found that the shortest intron in *T. vaginalis*, at only 23 nucleotides in size, is one of the shortest introns known to date. We tested our pipeline on a chromosome-scale assembly of the bird parasite *Trichomonas stableri*, the closest known relative to *T. vaginalis*. Our results revealed some conservation of the main features (total intron count, sequence, length distribution, and motifs) of these two closely related species, although differences in their functional annotation and duplication suggest more specialized splicing machinery in *T. vaginalis*.

## Introduction

*Trichomonas vaginalis* causes trichomoniasis, the most prevalent nonviral sexually transmitted infection (STI) worldwide, with an incidence of 354 million in 2019 [1]. Atypically for an STI, its prevalence is higher in women >40-50 years old, but it can produce adverse outcomes during pregnancy, including preterm birth, premature rupture of membranes, and low birth weight babies. Of major concern is that *T. vaginalis* infection is associated with a 1.5-fold increased risk for HIV acquisition [2, 3]. Despite its high morbidity and adverse clinical outcomes, trichomoniasis is considered a neglected disease since public awareness of it is low, research funding is scarce, and knowledge of the parasite at the molecular level is limited [4]. *T. vaginalis* is closely related to *Trichomonas* species found in the oral cavities of American columbid birds (pigeons and doves)[5, 6], supporting an ancestral origin of the species in this monophyletic order. The ancestor of *T. vaginalis* likely jumped from columbids to humans during a recent Holocene epoch spillover event following colombid colonization of the Americas, followed by adaptation of the parasite to the human reproductive tract [7].

The first draft genome of *T. vaginalis* strain G3, released by our group in 2007, consisted of ∼17,000 scaffolds comprising a total of ∼160 Mb and ∼60,000 protein-coding genes [8]. Among the key genomic characteristics educed from this assembly were the unusually large genome size for a unicellular eukaryote, a high proportion of repetitive DNA content (∼65%) consisting mainly of transposable elements (TEs), several massively amplified gene families, and a striking lack of introns (non-coding sequences within open reading frames). Only 65 *T. vaginalis* genes were predicted to contain one or more introns.

Initially assigned to “junk” DNA, spliceosomal introns play an important role in the molecular biology of eukaryotic cells. In addition to being involved in gene regulation and generating protein diversity, the functions of introns are usually divided into three main categories: (1) those associated with splicing, (2) non-coding DNAs (and related functions), and (3) storage of regulatory elements (including nested genes)[9–11]. It has been hypothesized that the origin and evolution of introns are closely associated with the evolution of genomes, making understanding introns essential to understanding the evolution of eukaryotes [9, 10].

The count of introns in *T. vaginalis* has varied widely over the years. In 2005, Vanacová *et al.* [12] experimentally confirmed the existence of functional splicing machinery in *T. vaginalis*. They identified single putative spliceosomal introns in 41 genes in a prepublication release of the *T. vaginalis* G3 assembly, with conserved motifs at both 5’ and 3’ ends of the introns, including canonical GT/AG and branch site (BS) similar to those needed for intron recognition by the splicing machinery in yeast and metazoa. The 65 intron-containing genes subsequently reported in the 2007 G3 draft genome sequence (5), were revised to 61 upon deposition to TrichDB (24). In 2009, Deng *et al.* [13] identified a gene with a new 25 bp intron, the shortest known, whose motifs differed from those previously described, suggesting the possibility of a more extensive intron repertoire. In 2018, Wang *et al.* [11] revisited the 62 putative intron-containing genes, using comparative PCR of their genomic DNA and cDNA to test each for splicing. They confirmed spliced introns in 31 of those genes, noted RNA-seq evidence for splicing in five more of them, found a new short intron in the 5’ UTR of a gene not on the list, and classified *T. vaginalis* introns into two types based on the conservation of the sequence and its length. The disparity between the initial conserved motif-based estimate and the number from experimental PCR testing was striking and given the divergent nature of the organism and the fragmented state of the draft genome assembly, it seemed likely that a definitive accounting of introns in *T. vaginalis* remained to be done.

We present here the most comprehensive analysis to date of the repertoire of spliceosomal introns in *T. vaginalis* G3, leveraging our new long-read chromosome-scale genome assembly, new RNAseq datasets, and a custom pipeline to identify genes containing introns and validating them using exon-intron boundary RT-PCR. We furthermore applied the same pipeline and analyses to a similarly high-quality chromosome-scale genome assembly from the closely related avian parasite *Trichomonas stableri* strain BTPI-3. Our results establish important similarities and differences between these two species and highlight specific features with potential impact on the evolution of trichomonads. We confirmed that trichomonad introns can be classified into two groups based on their length and sequence, identified conserved and extended intron motifs (including the BS), shortened the minimum intron length, expanded the repertoire of introns located in UTRs, and, for the first time, confirmed the existence of genes containing two introns.

## Results

### Custom, semi-automated pipeline for intron identification and curation

An initial mapping of our *T. vaginalis* strain G3 RNA-seq reads, generated from biological triplicate libraries as part of our project to generate a new chromosome-grade assembly of the parasite [14], suggested far more numerous splice events (or splice junctions (SJs)) than previously reported. Approximately 220 potential introns were identified (data not shown), including some that were unlikely >100 kilobases in length. Most of those SJs contained the universal and canonical GT/AG dinucleotides and were conserved among replicates, making it difficult to resolve whether they were real or artifactual. However, most of the questionable SJs were observed in regions enriched with TEs and containing significant GC troughs/peaks, indicating that those RNA-seq read alignments may have resulted from the highly repetitive content of the genome. A manual inspection of the predicted introns, combining a search for the motifs needed by the spliceosomal machinery with transcript assembly, identified at least 51 high-confidence intron-containing genes and enabled a recalculation of the consensus sequences of the motifs at both splice sites. The results also indicated that more introns might be present and the need for a more exhaustive analysis. Therefore, we designed an automated pipeline workflow to identify splicing events from RNAseq alignments by using the SJ coordinates and their flanking regions to search for conserved splicing motifs.

The semi-automated pipeline was developed to identify and assemble intron-containing genes in *T. vaginalis* using RNA-seq data and the degenerate intron motifs at the 5’ (GYAYGY, GTWYWD) and 3’ (RCTAACACAYAG, TCTAACH[1–2]AACAG) ends identified and confirmed by Vanacova *et al.* [12], Deng *et al* [13] and Wang *et al* [11]. The pipeline is summarized in **Fig 1**. Briefly, in the first step (Fig 1, blue), the RNAseq reads are mapped to the reference genome. In the second step (Fig 1, cyan), SJs mapping to regions lacking degenerate intron motifs are filtered out. Finally, transcript assembly is performed in the third step (Fig 1, yellow).

**Fig 1.**
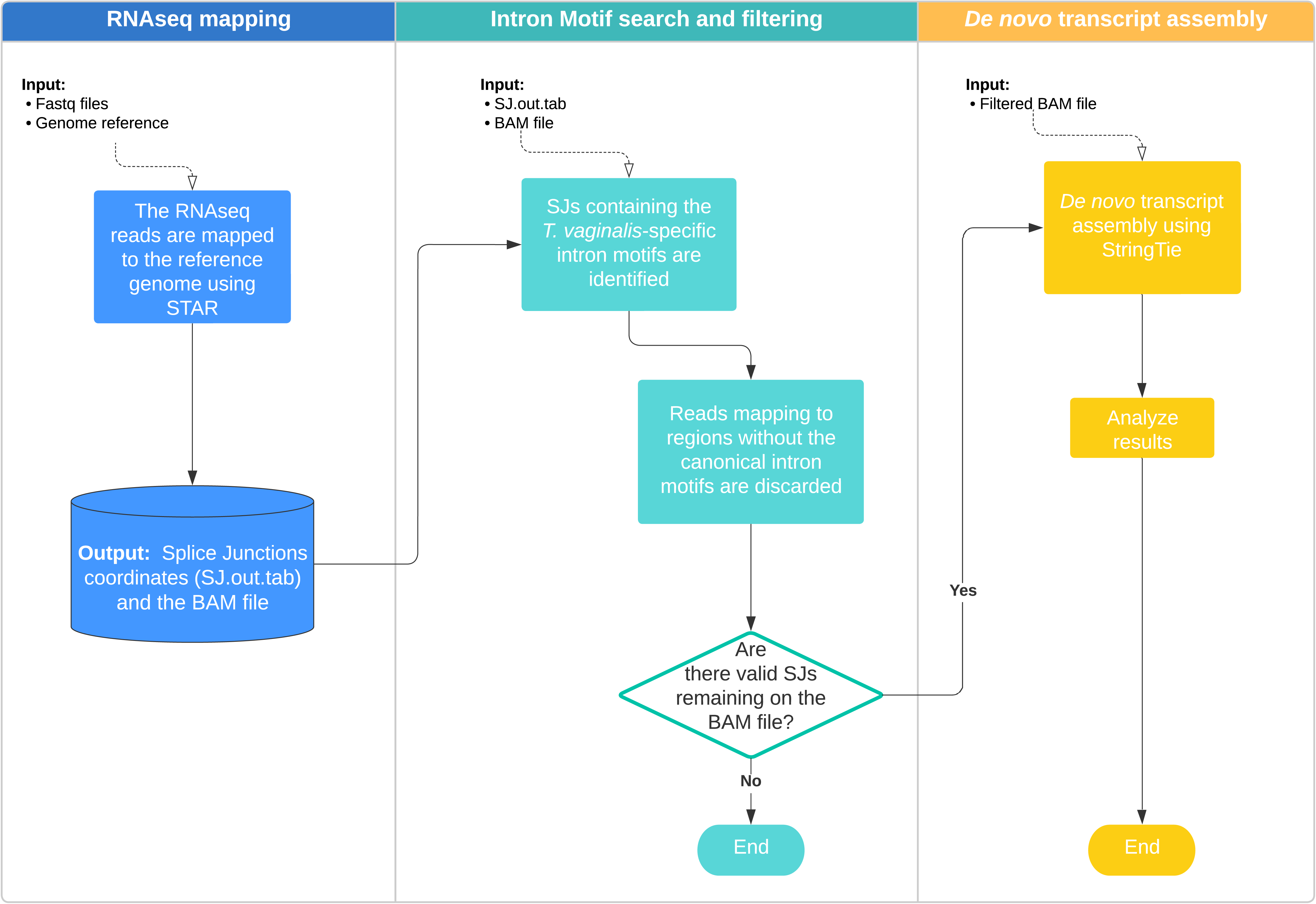
Semi-automated pipeline for the identification and assembly of intron-containing genes in *T. vaginalis* using RNAseq and degenerate intron motifs. In the first step, the RNAseq reads are mapped to the reference genome using STAR using default parameters (the mapping output file must be in a sorted BAM format and contain all SAM attributes). In the second step, SJs mapping to regions without the canonical intron motifs (degenerated intron motifs) are discarded. Finally, the transcript assembly is performed using the filtered version of the BAM file and StringTie using default parameters.

### Refinement of the *T. vaginalis* intron count

We ran the new chromosome-scale *T. vaginalis* genome sequence and RNAseq datasets in triplicate through our semi-automated pipeline. A total of 99.7% of the questionable SJs were filtered out, and the alignments passing the filter showed a clear pattern of introns with lengths of between ∼20 and ∼200 nucleotides (**S1 Fig**) in accord with the minima and maxima previously described for this parasite [11]. The list of putative intron-containing genes and their IDs can be found in **Supplementary Table 2** (**S2 Table**). After transcript assembly using StringTie, we identified single introns in 63 transcripts. Just over half (n=35) can be considered new since no earlier references to them were found, while 28 correspond to introns previously described by Vanacova *et al.* [12], Deng *et al.* [13], the 2007 draft genome annotation [8], and Wang *et al.* [11] (**Fig 2**).

**Fig 2.**
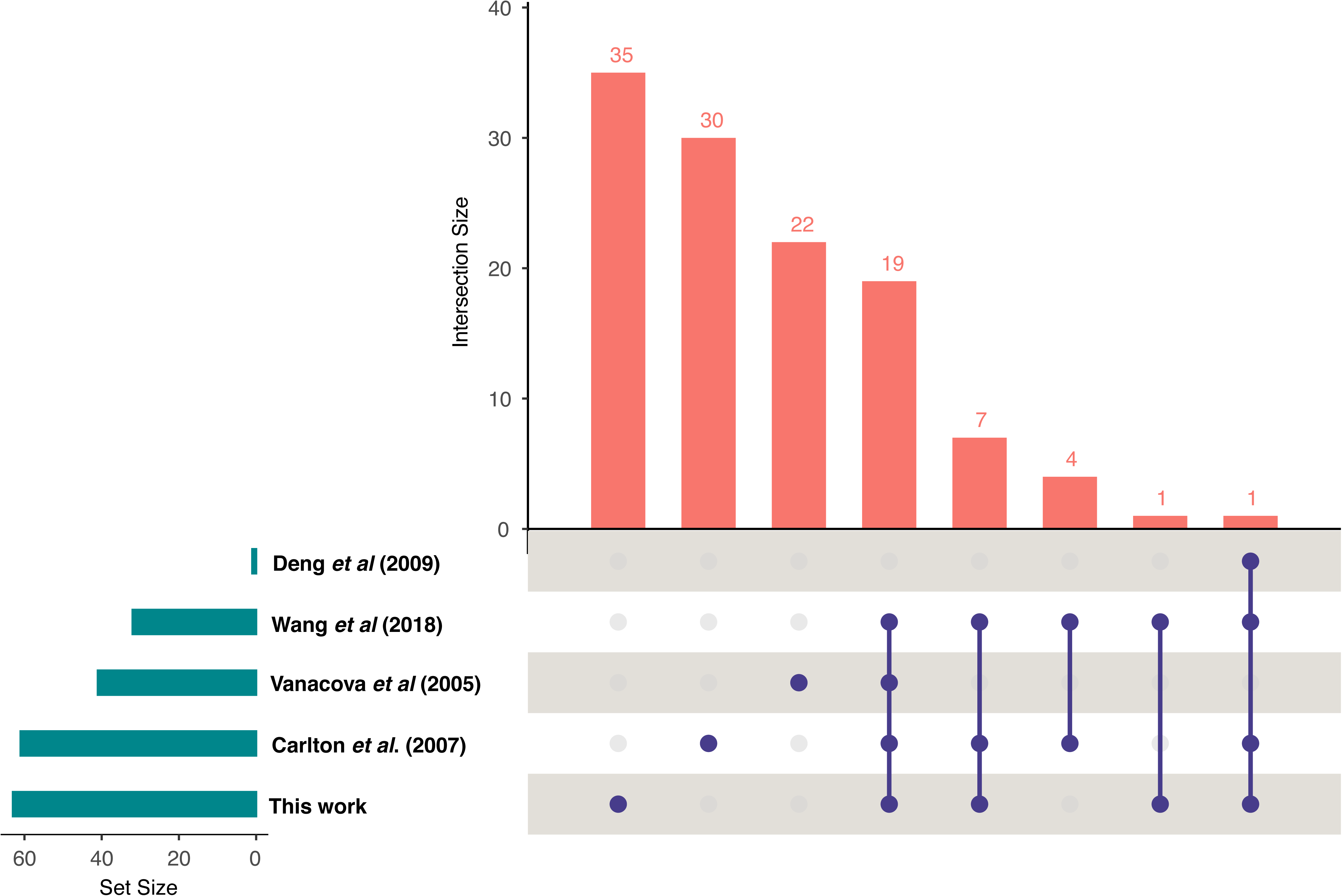
An upset plot summarizing the intron content in *T. vaginalis* G3 described by different authors since 2005 and the present work. The horizontal bars (dark cyan) indicate the total number of introns found by each author. Next to them, the intersection matrix shows the overlaps and unique elements among the different datasets. Finally, the height of the vertical bars (pink) indicates the number of elements in each intersection.

Four genes reported to have ‘functional introns’ by Wang *et al.* (2018) (TVAG_056030, TVAG_134480, TVAG_324910, TVAG_416520) were not validated by our results. Comparison of the primer sequences for TVAG_134480 to the new *T. vaginalis* G3 assembly shows that they match equally well to TVAG_089630, a 97% identical paralog containing a 26 bp intron. Therefore, we consider the Wang *et al.* PCR product for TVAG_134480 is a false positive -- an amplification of the TVAG_089630 intron. Wang *et al*. report the three other positives (TVAG_056030, TVAG_324910, TVAG_416520) that we could not validate as having a short (25-26 nt) intron length and classification as type B introns. We identified the CDS of these three genes in the new assembly (**S2 Fig**), but no introns were detected in them, and as we identified no paralogs or potential misassembles, we considered them false positives and excluded them from further analysis.

### Experimental validation of predicted *T. vaginalis* introns

We experimentally validated the 35 new intron-containing transcripts identified through our pipeline by PCR comparison of the genomic DNA (un-spliced) and cDNA (spliced) amplicon lengths (**Fig 3**). Transcript TVAGG3_0930320 is a 100% conserved paralog of TVAGG3_0931505 (including the intron, but on the opposite strand), and it was, therefore, not included in the figure. In addition, the transcript TVAGG3_0998030 is a paralog of the TVAGG3_0105790 gene (including the intron and both coding on the reverse strand) and was also not included in the figure. In agreement with previous results [8, 11, 12], most cDNAs produced both the un-spliced and spliced versions of the transcript, which was also confirmed by RNAseq reads crossing the intron sequence (**Fig 3**). RNAseq coverage results for our complete set of 63 intron-containing genes are shown in **Supplementary** Figure 3 (**S3 Fig**).

**Fig 3.**
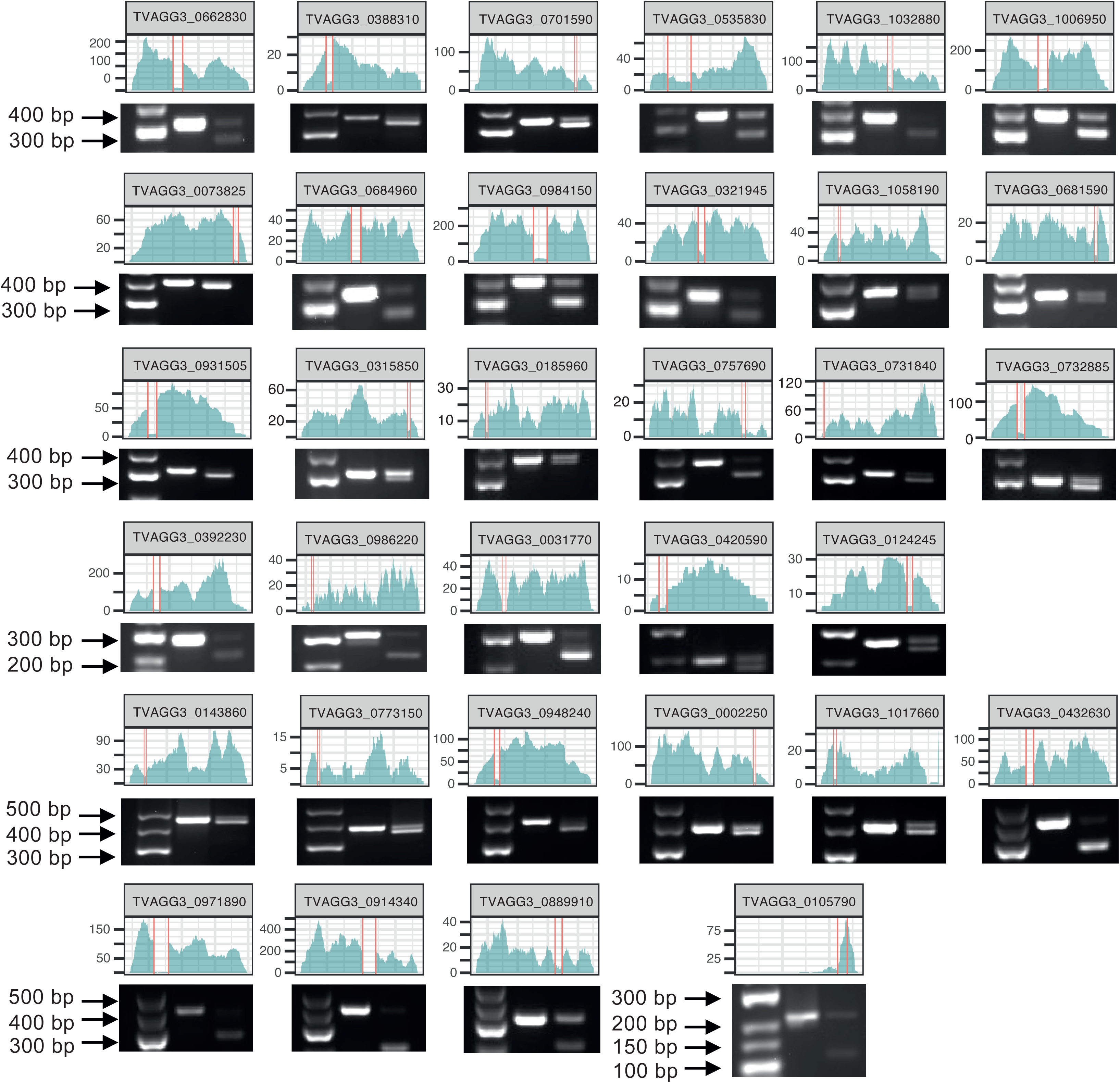
The 35 new intron-containing transcripts found in the *T. vaginalis G3* genome. The new introns (delimited by vertical red lines) were identified by RNAseq (coverage shown in cyan) and validated by PCR (lane 1; low range ladder, lane 2; gDNA product and lane 3; cDNA product). The paralogs TVAGG3_0930320 and TVAGG3_0998030 were not included in this figure.

In contrast with the results described by Vanacova *et al.* (2005), our findings confirmed that the 63 spliceosomal introns contain the GT/AG universal motifs at the splice site. Moreover, these intron signatures at both ends include an extended and highly conserved 5’ motif (up to seven nucleotides in total) and a BS (branch-site; ACTAA) fused with the 3’ motif for 12 nucleotides (**Fig 4**).

**Fig 4.**
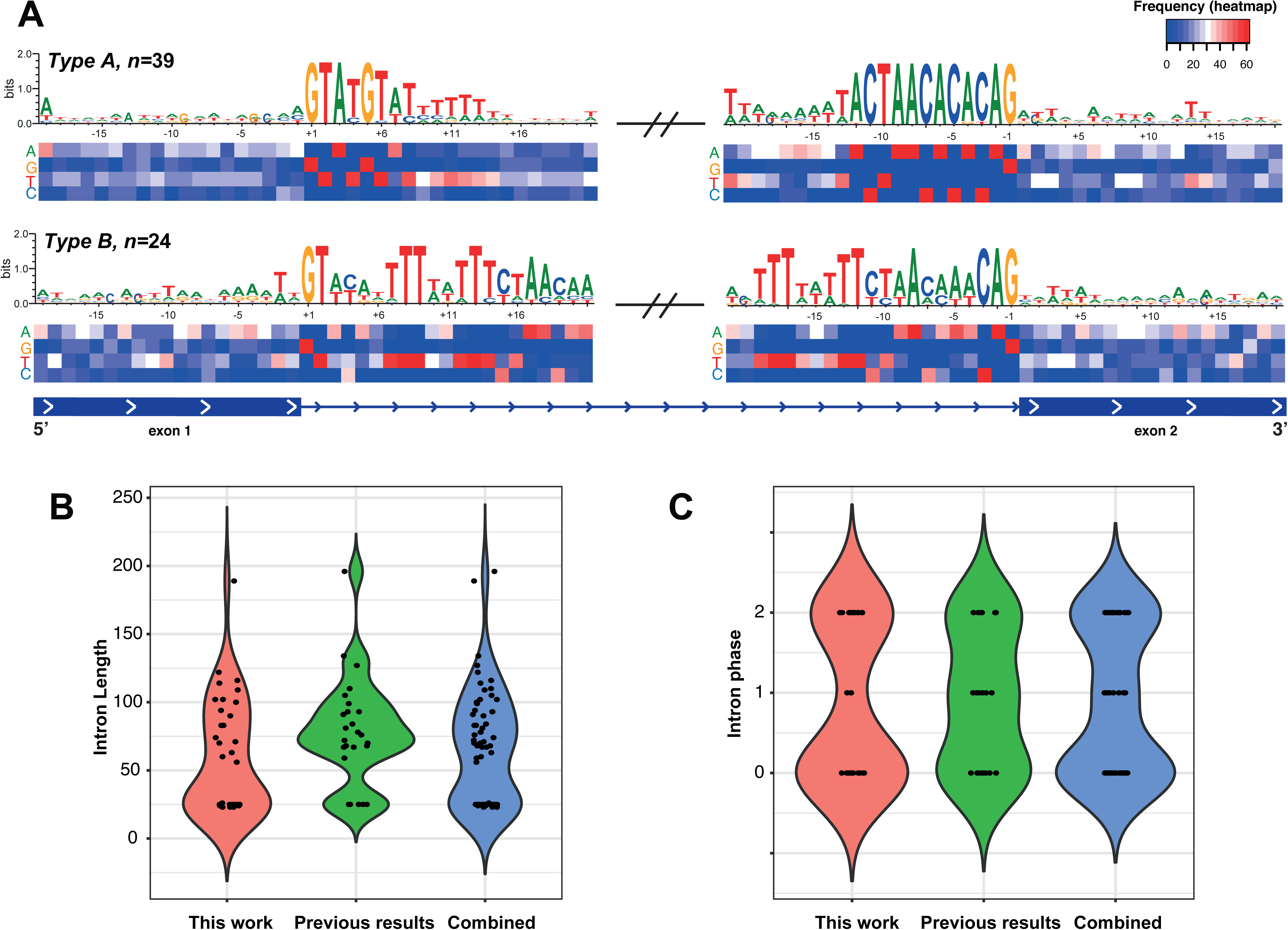
Consensus sequence motif at the splice sites and intron features in *T. vaginalis*. (A) The consensus sequence surrounding the exon-intron junction (+/-20 bp) for the 63 intron-containing genes is shown; the heatmap shows the nucleotide frequency at each position; (B) length distribution of the new introns compared to those previously described and the complete repertoire; (C) the intron phase distribution (excluding the 9 UTR introns). In both cases (panels B and C), the violin plots display the distribution of the values, and the plot width represents the density of data points at that value.

Wang *et al.* (2018) proposed classifying *T. vaginalis* introns into two types (A and B) based on conservation of intron motifs and their lengths. We found four out of the eight introns classified as type B in their results to be false positives. However, with the new repertoire including 35 novel introns, inferring two types of introns is still possible if we solely consider these two characteristics. *T. vaginalis* introns are thus either uniformly short (∼25 nucleotides, type B) or long (50-196 nucleotides, type A).

Twenty-four introns can be classified as type B. Of these, eighteen are introns newly described in our work, and six represent the shortest intron lengths reported for this parasite to date, with three sequences measuring 24 nt and three measuring 23 nt in length. We confirm that type B introns are characterized by a higher sequence variability at their 5’ and 3’ ends, including the BS and the consensus motifs (**Fig 4A**). The 5’ end is the more divergent; while positions +3 and +4 are composed of only two nucleotides (A/T and C/T), positions +5, +6, and +7 exhibit greater heterogeneity with up to three or four possible combinations. Positions +8 and +9 are well conserved with thymines. The 3’ end motif is comparatively variable. The first adenine of the BS motif (ACTAA) spanning from position −8 to −12 is replaced by thymine and followed by CTAA nucleotides with a single exception (TVAGG3_0315850) at position −11, where the cytosine is replaced by a thymine, in agreement with previous findings (Fig 4A and S4 Fig). In addition, it is also possible to identify a deletion at position −4 in the six shortest introns (S4 Fig). In both cases of the type B introns, the remaining sequence mainly comprises polypyrimidine tracts.

In contrast, type A introns (39 in total) are more variable in length, ranging from 56 up to 196 nucleotides, but with higher conservation of both motifs surrounding the splice site (**Fig 4 A, B**). The seven nucleotides of the 5’ end are well conserved with minor exceptions at positions +4, +6, and +7, and the 12 nucleotides of the 3’ end are almost perfectly conserved with minor exceptions at positions −12, −3 and −4. The first nucleotide of the BS (position −12) is predominantly composed of adenines rather than thymines, which agrees with previous findings. In contrast to type B introns, the polypyrimidine tract next to the 5’ and 3’ motifs is only slightly conserved at the 3’ end. The surrounding nucleotides of both motifs exhibit a significantly lower percentage of G+C compared to the exon sequences, and no additional consensus motifs were identified.

Although most introns in *T. vaginalis* (and other eukaryotes) can be classified as phase zero, one, or two (**Fig 4C**), Wang *et al.* reported the presence of one intron located at the 5’ UTR end of the TVAG_269270 gene that does not follow such classification. Our findings confirmed this result and produced a significant expansion of this class of introns (referred to as “UTR introns”), identifying nine new instances with only one in the 3’ UTR (TVAGG3_0535830, **S2 Table**). Despite the classification of most (7/10) of these UTR introns as type B, no additional shared characteristics, such as functional annotation or sequence conservation, were identified. In line with observations in many other organisms, phase 0 introns are the most abundant in this parasite, totaling 25 instances, followed by 19 phase 2, and the phase 1 introns are less represented, with only nine occurrences.

The predicted functions of the proteins encoded by the intron-containing genes are equally diverse, ranging across protein phosphorylation, transcription, intracellular signaling, cellular movement, and ‘hypothetical’ (unknown function), among others (**Table 1**). Most functions are found in multi-gene families (*e.g.,* kinases, Armadillo-type repeats, leucine-rich, dyneins, cyclins, and GTPases). The most common products are those with predicted kinase functions. All of their introns are type A, but the motifs at both splice sites are not necessarily conserved (especially at the 5’), with phase 0 being the most frequent feature.

**Table 1.** Functional annotation of the intron-containing genes in *T. vaginalis* and intron attributes.

| Product name | Count | Gene IDs | Intron type | Intron phase |
| --- | --- | --- | --- | --- |
| Protein kinase domain | 16 | TVAGG3_0860620, TVAGG3_0764700, TVAGG3_0199570,<br>TVAGG3_0661130, TVAGG3_1058000, TVAGG3_1043790, | A | 0:11, 1:5 |
|  |  | TVAGG3_0677070, TVAGG3_0972410, TVAGG3_0299880,<br>TVAGG3_0511450, TVAGG3_0403570, TVAGG3_0034870,<br>TVAGG3_0914340, TVAGG3_0971890, TVAGG3_0321945,<br>TVAGG3_0392230 |  |  |
| Unknown | 11 | TVAGG3_1019300, TVAGG3_0002250, TVAGG3_0124245,<br>TVAGG3_0315850, TVAGG3_0388310, TVAGG3_0732885,<br>TVAGG3_0143860, TVAGG3_0185960, TVAGG3_0986220,<br>TVAGG3_0757690, TVAGG3_0773150 | A:3, B:8 | 0:3, 1:2, 2:4,<br>5UTR:2 |
| Armadillo-type fold | 6 | TVAGG3_0108840, TVAGG3_0283200, TVAGG3_0841800,<br>TVAGG3_1058190, TVAGG3_0701590, TVAGG3_0681590 | B | 2:4, 5UTR:2 |
| FCP1 homology domain | 3 | TVAGG3_0288310, TVAGG3_0182080, TVAGG3_0644560 | A | 0 |
| Initiator binding domain | 3 | TVAGG3_0984150, TVAGG3_1006950, TVAGG3_0662830 | A | 2 |
| Ankyrin repeat-<br>containing domain | 3 | TVAGG3_0031770, TVAGG3_0136820, TVAGG3_1017660 | A:2, B:1 | 0:1, 2:1,<br>5UTR:1 |
| Dynein light chain<br>Tctex-1 like | 2 | TVAGG3_0930320, TVAGG3_0931505 | B | 5UTR |
| Arf GTPase activating<br>protein | 2 | TVAGG3_0432630, TVAGG3_0889910 | B | 0 |
| Pleckstrin homology<br>domain | 2 | TVAGG3_0656270, TVAGG3_0997450 | A, B | 2, 0 |
| Small GTPase | 2 | TVAGG3_0006570, TVAGG3_0073825 | B | 2 |
| Pre-mRNA-splicing<br>factor 18 | 2 | TVAGG3_0105790, TVAGG3_0998030 | A | 5UTR |
| Cyclin Dependent<br>Kinase Inhibitor 2C | 1 | TVAGG3_0948240 | B | 0 |
| Leucine-rich repeat<br>domain superfamily | 1 | TVAGG3_0731840 | B | 0 |
| LSM domain<br>superfamily | 1 | TVAGG3_0420590 | B | 0 |
| MOB kinase activator<br>family | 1 | TVAGG3_0032670 | B | 2 |
| P-loop containing<br>nucleoside triphosphate<br>hydrolase | 1 | TVAGG3_1032880 | A | 0 |
| Palmitoyltransferase,<br>DHHC domain | 1 | TVAGG3_0684960 | A | 2 |
| Polymerase, nucleotidyl transferase domain | 1 | TVAGG3_0321030 | A | 1 |
| RNA-Binding ASCH domain | 1 | TVAGG3_0998660 | A | 1 |
| TAF6, C-terminal HEAT repeat domain | 1 | TVAGG3_0852200 | A | 2 |
| Transcription initiation factor TFIID subunit 6 | 1 | TVAGG3_0998160 | A | 2 |
| U6 snRNA-associated Sm-like protein Lsm3 | 1 | TVAGG3_0535830 | A | 3UTR |

Eleven genes were classified as encoding hypothetical proteins (or proteins with unknown function), the second most frequent annotation. This group’s introns are more divergent than those from the protein kinases, containing both type A and B introns in all three phases and having two UTR introns.

The third most frequent “function” comprises proteins containing Armadillo repeats. These genes are also characterized by short intron length (type B), phase 2, and two 5’UTR introns. Other genes with repetitive domains (ankyrin and leucine-rich, mainly) showed higher divergence, with both types of introns and phase 0, phase 2, and 5’ UTR introns. In all cases, no correlation was found between the functional annotation and the conservation of the intron motifs.

Interestingly, three intron-containing genes encode proteins with predicted roles in the spliceosomal machinery (U6 snRNA-associated protein and pre-mRNA splicing factor), two of which are almost perfectly conserved paralogs (TVAGG3_0105790, TVAGG3_0998030) and the other being the only one with an intron located in the 3’ UTR (TVAGG3_0535830). The three introns are type A (100 and 102 nucleotides in length for the two paralogs and TVAGG3_0535830, respectively).

### Intron prediction and validation in the closest relative of birds, *T. stableri*

To validate our results and pipeline, we conducted the same analysis for the sister species of *T. vaginalis*, the bird parasite *Trichomonas stableri* (strain BTPI-3). This trichomonad infects columbid birds (pigeons and doves) and was associated with widespread Pacific Coast band-tailed pigeon mortality in California in 2006–2007 [15]. We analyzed the intron differences and similarities between these two species using the same pipeline described in **Fig 1** and a new chromosome-scale *T. stableri* BTPI-3 genome assembly and triplicate RNA-seq dataset generated by our group [14]. We identified a total of 81 *T. stableri* intron-containing genes (versus 63 in *T. vaginalis* G3), 78 with one intron per gene, and for the first time in a trichomonad, three genes with two introns.

We found an unexpected degree of divergence between these two species in their intron characteristics: first, in the total number of intron-containing genes, second, in the number of introns per gene, and third, in the conservation of the intron motifs (**Fig 5**). *T. stableri* BTPI-3 type A introns are similarly abundant to *T. vaginalis* (32 versus 39) and have the same length distribution (56-196 nucleotides). Type B introns are more than twice as abundant in *T. stableri* (52 versus 24) compared to *T. vaginalis*, with a length varying from 23-28 nucleotides (23-25 in *T. vaginalis*), and a higher variability in the consensus intron motifs (**Fig 5**). Notably, the introns present in the three transcripts with three exons are all type B (**Fig 5, A-B**), and we found polymorphisms differing from the *T. vaginalis* G3 consensus motifs in one of them (TsBTPI-3-004276), in both the 5’ and 3’ ends. In the 3’ end, we found SNPs at positions −3 (A), −8 (C), −10 (A), and −12 (C), and one SNP in the 5’ end at position +4 (A) (**Fig 5A**). Given these polymorphisms, the DNA and mRNA sequences for this gene were validated by PCR (**Fig 5C**). In summary, *T. stableri* BTPI3 exhibits a wider and more variable sequence repertoire of short introns (with up to two introns per gene) and a conserved repertoire of large introns (type A), compared to *T. vaginalis* G3.

**Fig 5.**
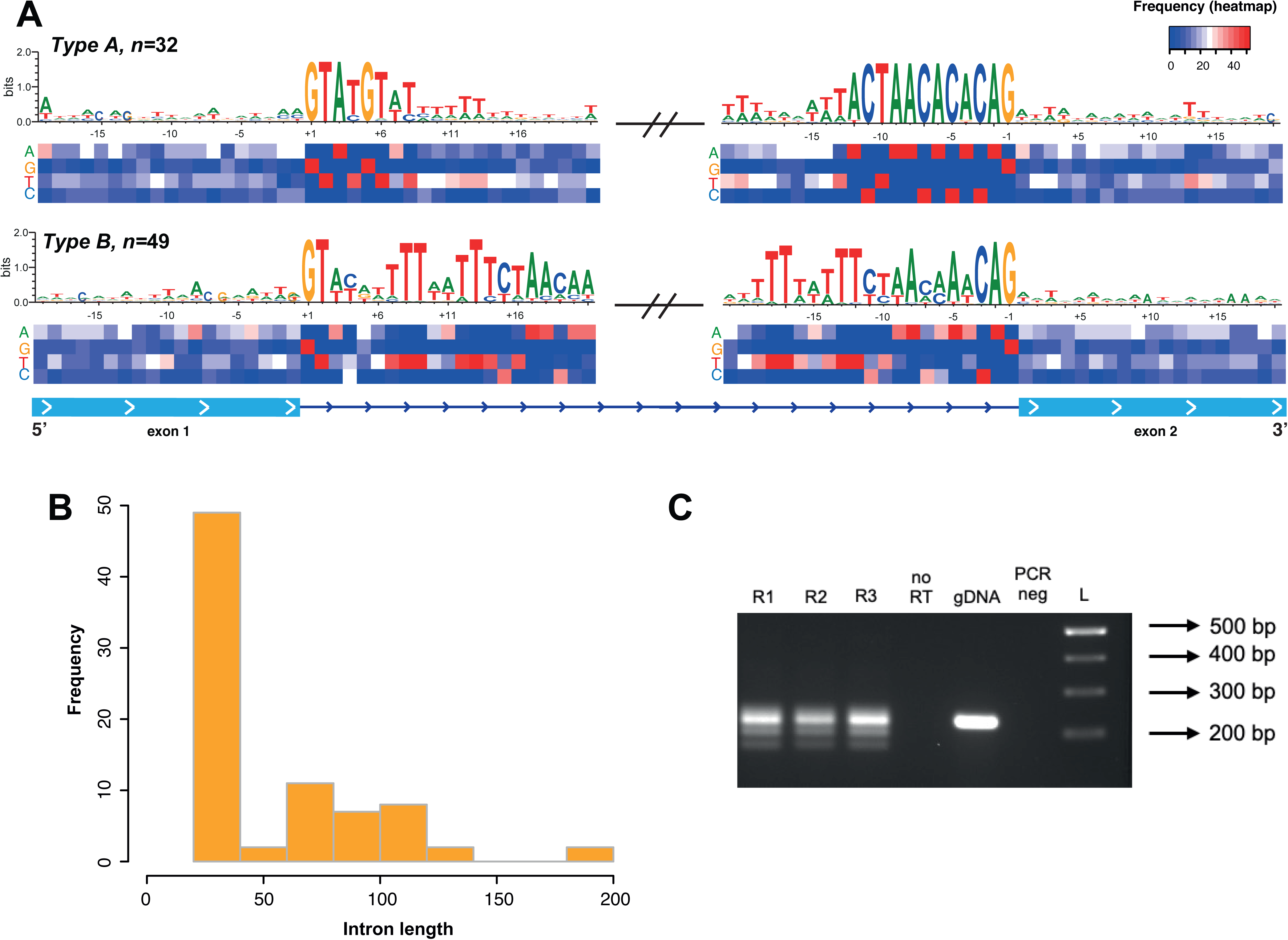
*T. stableri* BTPI-3 intron features. Consensus sequence motifs and nucleotide frequency are shown in panel **A**, an histogtram of the intron length distribution is shown in panel **B**, and the PCR validation of the gene TsBTPI-3_004276 in panel **C**; lanes 1,2, and 3 are the replicates of the cDNA amplification, lane 4; RNA, lane 5 gDNA amplification, lane 6; negative control, and lane 7 the 1k Plus DNA ladder.

Of the 81 intron-containing genes identified in *T. stableri* BTPI3, 50 (∼62%) are conserved in *T. vaginalis* G3 with an average identity of 91% (+/-3%, including the intron sequence). However, we only found signatures of positive selection (dN/dS ratio > 1) in 24 of these genes (S2 table), and their functional annotations suggest they might be involved in constitutive functions such as gene expression, protein trafficking, cellular movement, and energy production. Interestingly, for 11 of these conserved genes, the spliceosomal intron is perfectly conserved, and all of them are classified as type B introns (length 23-26 nucleotides). The remaining 39 conserved genes exhibited variable intron sequences between the species, although we did not detect a significant change in their length that would classify them into different groups (A or B) or affect the intron phase among species. Most of the conserved and species-specific intron-containing genes were predicted to have similar functions in the two species or belong to the expanded gene families (S2 table).

The intron phase distribution did not show significant differences between the two species, *e.g.,* phase 0 introns were also confirmed as the most abundant in *T. stableri* BTPI*-3* (30 introns versus 25), and similar numbers for phase 2 (21 vs 19 in G3) and UTR introns (ten in both species) were found. In contrast, only phase 1 introns exhibited significant expansion, with 20 instances versus 9 in *T. vaginalis*.

Finally, we performed a *de novo* search in both species’ genomes for intron sequences not captured by RNAseq and/or the transcript assembly but containing the newly identified putative intron motifs (5’ and 3’). The search was performed on both strands, no overlapping or redundant introns were allowed, and only potential introns with a maximum length of 200 nucleotides were considered. Our search yielded 34 potential introns in the *T. vaginalis* G3 genome (20 type A and 14 type B). Of those, only nine are located within or near an annotated ORF (**S3 Table**). In *T. stableri* BTPI3, we identified a total of 20 likely valid sequences (**S3 Table**). The majority were classified as type A (19), with only three sequences as type B, and nine of those are located within or close to an annotated ORF.

## Discussion

Due to the sequence repetitiveness of the *T. vaginalis* genome (largely from multiple copies of TE families and multicopy protein-coding gene families), standard RNAseq mapping reported thousands of likely artifactual chimeric splice junctions, suggesting hundreds of spliced transcripts, many spanning distances ranging from few kilobases to hundreds of thousands of kilobases. Although such mapping issues are well known in repetitive sequences with few solutions to this problem available [16], motifs conserved in the 5’ and 3’ ends of the introns that are specific to the splicing machinery in *T. vaginalis* enabled us to identify and filter out artifactual mappings by referring to their genomic sequences. In line with this observation, recent analysis using a logistic regression classifier has shown conservation patterns on the splice sites across more than 400 species, confirming that the annotation of spliceosomal introns goes beyond the GT/AG dinucleotides and might lead to erroneous annotations [17]. Armed with this knowledge, we developed a custom pipeline to identify spliceosomal introns using RNAseq data, allowing us to finally identify the complete repertoire of intron-containing genes in *T. vaginalis* G3, and identify introns for the first time in its sister species, the bird parasite *T. stableri* BTPI-3. Our results significantly expanded the repertoire and knowledge about spliceosomal introns in these parasites, including the number of introns, intron size, and the consensus motifs.

A combination of the new chromosome-grade assembly of *T. vaginalis* G3 [14] -- constituting ∼180 Mb in six chromosome-length scaffolds and ∼44,000 putative protein-coding genes (plus ∼47,000 transposable element genes) -- and our results, identifying and validating 35 new spliceosomal introns for a total of 63, allows us to finally confirm a low per-kilobase intron density across the complete genome, and conclude that *T. vaginalis* is an intron-poor parasite. Furthermore, functional annotation of the new assembly indicates the existence of only 57 genes associated with the spliceosome machinery [14], supporting the hypothesis that intron-poor genomes are also constrained in their spliceosome machinery [18]. A chromosome-scale assembly of the closest sister species to *T. vaginalis,* the bird parasite*T. stableri* BTPI-3, generated by our group and composed of ∼70 Mb in six chromosomes and ∼32,000 putative protein-coding genes [14], and the results presented in this manuscript, further confirm that trichomonad genomes have a low intron density generally, while exhibiting interesting similarities and differences in their intron characteristics, e.g., a more extensive repertoire of introns in *T. stableri* BTPI-3 despite a shorter genome length and lesser gene count compared to *T. vaginalis* G3; conservation of intron motifs needed by the splicing machinery (including the BS); and a highly conserved intron length distribution.

Although our identified intron-containing genes code for proteins with unknown functions or belong to the highly expanded multicopy gene families, some exceptions suggest they are also involved in essential molecular mechanisms such as transcription (**Table 1 and S2 Table**). Duplication of some of these genes in the *T. vaginalis* genome might accord with this hypothesis and suggest a more sophisticated transcription control. Overall, our results describing the intron content and their most relevant features in two species of genus *Trichomonas* can be used to track the relevance of the splicing machinery during the evolution of this eukaryote and in comparison with other species.

We could not confirm four of the previously described introns in *T. vaginalis*, and we believe this is due to the fragmentation and misassembly of the 2007 draft genome. We found that one of those false positives and two newly validated introns (including the CDS) could be used to understand the activity of DNA transposons in the genome. First, we found that a true positive intron-containing gene (TVAG_089630) has an intron-less paralog (a false positive) in the opposite strand (previously named TVAG_134480), suggesting that it may have been duplicated by a transposable element (or a retrotransposon after the intron removal). Second, TVAGG3_0930320 and TVAGG3_0931505, both containing novel introns, are perfectly conserved paralogs, which also suggests their duplication by DNA transposons before intron removal. Our genome annotation results [14] also support this hypothesis, confirming that both pairs of paralogs are flanked by TEs. These results suggest that gene duplication through transposable element high-jacking in *T. vaginalis* is a highly selective and stringent process, given the massive expansion of transposable elements that has occurred within the genome and the very low intron density. In agreement with this hypothesis, we did not identify intron-containing genes undergoing these mechanisms in *T. stableri* BTPI-3. On the other hand, a third pair of intron-containing *T. vaginalis* gene paralogs located on different chromosomes but the same strand (TVAGG3_0105790 and TVAGG3_0998030) suggest that their duplication was produced using an alternative mechanism since they are not flanked by TEs in contrast to the former two pairs of paralogs.

The shortest intron we identified in both *Trichomonas* species is 23 nucleotides long, among the shortest known and comparable only with the 15 nucleotides of an intron in the *Stentor coeruleus* (trumpet ciliate) genome [19] and the 18 nucleotide intron in *Bigelowiella natans* (unicellular marine algae) genome [20]. The longest intron we confirmed is 196 nucleotides in *T. vaginalis*, which is short compared with most eukaryotes, where maximum intron length varies from a few hundred bases to tens of kilobases [21, 22]. While it conforms with the hypothesis that intron size is positively correlated with genome size, where short genomes (<500 Mb) tend to have short introns (<1 kb) and less specialized spliceosomal machinery [10, 18], our results are in contrast to the suggestion that short introns (less than 40 bp) cannot form complete introns and do not contain regulatory elements [9, 19, 23]. We found that the simplicity of the splicing machinery corresponds with the simplicity of the spliceosomal repertoire, including short introns, the absence of multiple introns per transcript, the absence of potential isoforms, and the absence of alternative splicing insertion sites. Besides, our results in both species of trichomonads confirm the conservation of motifs (including the BS) needed by the splicing machinery.

We also confirmed that introns in *Trichomonas* species could be classified into two groups based on their length and intron motifs, with type A comprising longer introns (>50 nt) and type B comprising shorter (<<50 nt) ones, as suggested by Wang *et al*. [11]. Studies in diplomonad and parabasalid genomes have demonstrated that these differences in intron length do not affect the processing efficiency of the spliceosome, for example, through the formation of stem-loops and shortening the distance between exons and the branch site with the splice sites, facilitating intron removal independent of their length [18].

An additional interesting feature of *T. vaginalis* introns is the presence of extended consensus motifs at both splice sites (**Fig 4A**). Since their discovery, the high similarity to these present in *Giardia* and *Saccharomyces* has been demonstrated [12, 24]. However, in *T. vaginalis,* the 5’ motif is extended up to seven nucleotides, and the 3’ motif is fused with the BS for a total of 12 nucleotides. This contrasts with most eukaryotes (with a longer length distribution), where the BS is separated from the 3’ splice site by a non-conserved sequence of variable length (usually a polypyrimidine tract)[24]. The conservation of this structure in this deep-branching eukaryote might confirm that the efficiency of the splicing machinery relies on the conservation of the splice site and the BS, while the capacity of the intron sequence to form secondary structures to shorten the distance between the BS and the 3’ splice site is a complementary mechanism evolving differently between species [12, 18].

The intron phase is another feature used for intron classification and analysis of their evolution [9, 10]. It refers to the intron position relative to the codons of the flanking exon sequences (phase 0 falls between two codons, phase 1 between the first and second nucleotide, and phase 2 between the second and third nucleotide of the flanking codon). In most eukaryotes, where U2-type introns are the major spliceosome (as is the case of *T. vaginalis* and *T. stableri*), phase 0 introns are more abundant than phase 1 (the second most abundant) and phase 2 [9]. We found phase 0 introns to be the most abundant in both *Trichomonas* species (S2 Table). However, we found two interesting exceptions for phases 1 and 2 in *T. vaginalis*. First, phase 2 is the second most expanded (19 in total), and second, phase 1 is as abundant as those located in the UTRs (10 instances each). Furthermore, the high abundance of introns we found located in UTRs (nine in the 5’UTR and one in the 3’UTR in *T. vaginalis*) contrasts with their very low occurrence in most organisms [22, 25]. Second, the excess of phase 2 over phase 1 introns is typically found in U12-dependent introns [21], but U12 spliceosome subunit genes appear to be absent in the *T. vaginalis* [14]. Also, the strong conservation of intron motifs (including the BS conserved in *Giardia* and *Saccharomyces* specific to U2 introns [18, 24]) supports the hypothesis that U12-dependent introns are absent in *T. vaginalis* and *T. stableri*. Consequently, these results challenge the “intron-first” hypothesis, which posits that present-day introns derive from sequences between minigenes in the progenote and, therefore, must lie in phase 0. Instead, our results align more closely with the “introns-late” hypothesis, which suggests that the non-uniformity of intron phase distribution reflects the nonrandomness of intron insertions.

Finally, it is also important to note that the intron motifs located in UTRs are as well conserved as those found in the CDS, and no significant differences were found among them or when compared with CDS introns. Furthermore, similar numbers of this class of introns were validated to exist in both species. Our results accord with the assumption that introns in 3’UTRs are less abundant than those found in 5’UTRs, but more importantly, we attest that this class of introns should not be ignored or considered less relevant [26], especially in a low intron-density organism. For instance, it has been reported that introns located in 5’ UTRs contain regulatory elements such as small nucleolar RNA genes and microRNA precursors and, therefore, can be subjects of selection, playing an important role in gene expression [25, 27–29]. Further studies are needed to understand the biological relevance of this class of introns and their conservancy in these species over evolution.

Further analysis will also confirm if the additional intron sequences we detected by a *de novo* search (34 in *T. vaginalis* and 20 in *T. stableri,* but not validated by RNAseq) using the newly recalculated intron signatures are remains of old genes or if they are part of active genes expressed only in alternative and less common parasite forms, such as pseudocysts [30].

## Conclusions

We used a new chromosome-scale genome assembly, RNAseq data, and a custom pipeline to filter chimeric alignments to report the most complete repertoire of spliceosomal introns in the human pathogen *T. vaginalis*. We confirmed the existence of 35 new introns for a total of 63 (one per gene). These new results confirmed some of the most relevant features previously described using the draft genome, such as conserved motifs at the intron boundaries and their short length. But our results significantly expand information about intron features that can contribute to understanding the evolution of this parasite, including revising downwards the minimum intron length, establishing sequence conservation at the splice site at higher resolution, and describing the phase distribution with a significant increase of introns at the UTRs. We also confirmed the absence of U12-type introns, the ubiquitous presence of the GT/AG dinucleotides, the conservation of the BS fused with the 3’ splice site, and a few genes potentially duplicated by RNA and DNA transposons.

By applying the same methods to high-quality sequence data from a related species, we also generated the first repertoire of spliceosomal introns in the bird parasite *T. stableri* BTPI-3, consisting of 81 examples, of which three are genes containing two introns. Our results showed a high conservation of splicing signals between species. However, observed differences prompt further investigation to analyze the independent evolution of introns and their implications for the molecular biology of the parasites.

Both species of *Trichomonas* we studied appear to possess intron-poor genomes. The functional annotation of these genes suggests an enriched repertoire of constitutive functions, with *T. vaginalis* potentially harboring a more specialized spliceosomal machinery than *T. stableri*. On the other hand, an excess of phase zero introns and the conserved non-uniform distribution of phase one, phase two, and UTR introns in both species agree with observations in most eukaryotes of the introns-late hypothesis. Our results contribute to understanding introns in non-model organisms and their evolution over time.

## Materials and Methods

### Parasite culture and nucleic acid extraction

*T. vaginalis* strain G3 and *T. stableri* strain BTPI-3 parasites were grown in axenic culture at 37°C in modified Diamond’s Medium [31] supplemented with 10% horse serum, penicillin, and streptomycin (Invitrogen), and iron solution composed of ferrous ammonium sulfate and sulfosalicylic acid (Fisher Scientific), as described previously [32]. Cultures were grown overnight in 10 ml of media in 15 ml loose-capped conical tubes. Total DNA was extracted from *T. vaginalis* and *T. stableri* cultures using the DNeasy Blood and Tissue kit (Qiagen) following the manufacturer’s instructions. Total RNA from *T. vaginalis* G3 was extracted from biological triplicate cultures using the RNeasy Mini Kit (Qiagen) according to the manufacturer’s instructions. Total RNA from *T. stableri* BTPI-3 was extracted with TRIzol (Ambion #15996018) following the manufacturers’ instructions. DNA and RNA concentrations were determined using a Qubit fluorometer.

### *Trichomonas* genome and transcriptome datasets

Chromosome-grade assemblies of *T. vaginalis* strain G3 (PRJNA885811) and *T. stableri* strain BTPI-3 (PRJNA816543) were used. Transcriptome datasets consisting of RNA-seq reads from libraries of biological triplicates of *T. vaginalis* G3 (SRR22985603-SRR22985607) and *T. stableri* BTPI-3 (SRR22985597-SRR22985599) were also analyzed. The quality of the RNA-seq reads was assessed using FastQC [33] and were filtered and trimmed using Trimmomatic v0.39 [34] to a minimum length and overall quality of 70 nucleotides and 28 (phred score), respectively; adapter contamination was also removed when present.

### Semi-automated pipeline for intron validation

A semi-automated pipeline for intron validation was generated using a collection of bioinformatics tools and in-house python scripts. First the RNAseq reads are mapped to the reference genome using STAR v.2.7.6a [35], with no custom parameters except *--outSAMattributes All* to report all the alignment attributes in the alignment file and *--outSAMtype BAM SortedByCoordinate* to sort the BAM file by read coordinate. Second, SJs mapping to regions lacking degenerated intron motifs are filtered out from the BAM file using an in-house Python script and the SJ coordinates reported by STAR in the SJ.out.tab file. Third, the transcript assembly is carried out using StringTie (v.2.1.6, default parameters), and the resulting BAM file from step 2 (the BAM file must be sorted and indexed using Samtools[36]). The pipeline is available at https://github.com/biofcallejas/pysplicing. An additional search of potentially functional introns not captured by the RNAseq or previous analyses was performed using custom in-house Python scripts and the consensus sequences determined in the previous section. The search was performed in both strands; no overlaps were allowed and the maximum intron length was set to 200 nucleotides.

### Experimental validation of introns

Primer3 plus [37] was used to design pairs of PCR primers flanking predicted intron sequences. Their specificity was confirmed using a local version of Blastn (using *-task blastn-short*) and the new genome assemblies. The complete set of primer sequences is available in **supplementary table 1** (**S1 Table**). *T. vaginalis* G3 and *T. stableri* BTPI-3 cDNA were generated as follows: polyA mRNA was isolated from total RNA using the NEBNext Poly(A) mRNA Magnetic Isolation Module (#E7490), and cDNA was synthesized from poly(A) mRNA with oligo(dT) using the SuperScript III First-Strand Synthesis System (Invitrogen #18080051) following the manufacturer’s instructions. PCRs were performed using DreamTaq Green DNA Polymerase (Thermofisher) and standard conditions for 30 cycles. We used the previously validated (Wang *et al*. (8) and confirmed by our RNAseq results) TVAG_350500, and TVAG_306990 as positive controls. Negative controls were from cDNA synthesis reactions using poly(A)+ mRNA but no reverse transcriptase. PCR products were compared side by side on a 2.0% (w/v) agarose electrophoresis gel stained with 3 μl of ethidium bromide and a BioRad Low Range DNA Ladder.

### Functional annotation and intron features

Open reading frames for the intron-containing genes were predicted using Augustus [38] (local version v.3.5.0) and the standard genetic code. Functional annotation was predicted using the protein domain identifier InterProScan (v 5.50)[39]. Intron phase was determined using the structural annotation from the previous step and in-house Python scripts, considering the longest ORF crossing the exon-exon junction. Multiple sequence alignment (MSA) for all the validated splicing sites was calculated using MAFFT (BLOSUM62) [40] and visualized using Jalview [41].The genome and RNAseq alignments were visualized using IGV v.2.16.0 [42]. Consensus sequences were generated using WebLogo v2.8.2[43] and formatted using Adobe Illustrator. All other figures were generated using R and Rstudio [44].

## Acknowledgments

Research reported in this publication was supported by the National Institute Of Allergy And Infectious Diseases of the National Institutes of Health under Award Number R21AI149449. This work was supported in part through the NYU IT High-Performance Computing resources, services, and staff expertise.

## Supporting information

**S1 Fig.**
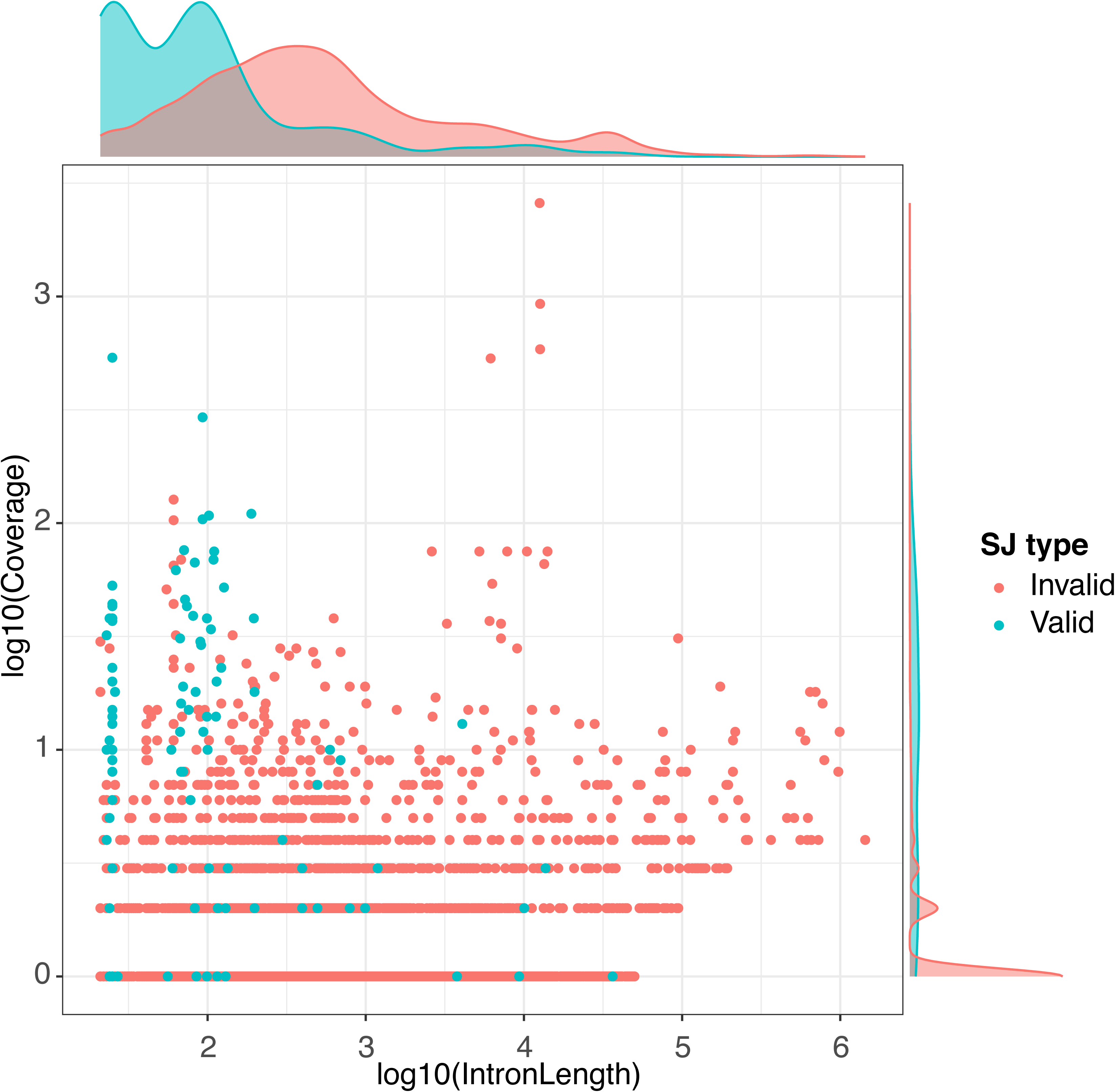
Splice junctions detected by mapping the RNAseq data to the new chromosome-scale assembly of *T. vaginalis G3*. Coverage and length (log10-based) for each SJ are shown in the “Y” and “X” axes, respectively. SJs were classified as invalid and valid by filtering those mapping to genomic regions lacking the degenerated intron motifs needed by the spliceosomal machinery.

**S2 Fig.**
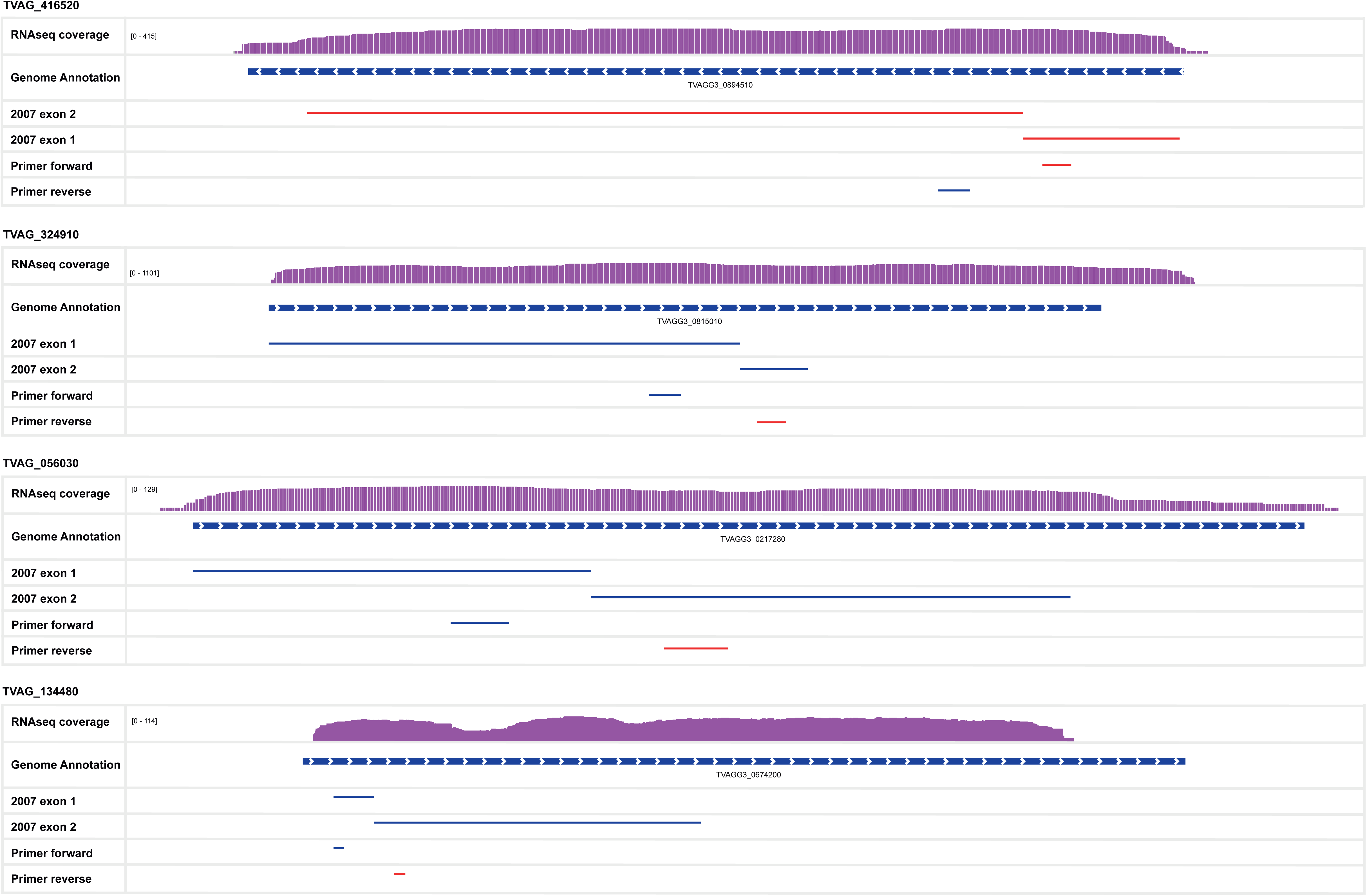
False positive intron-containing genes reported by Vanacova *et al.* (2005) and Wang *et al.* (2018). The RNAseq coverage (violet) mapped to the new genome and the new genome annotation (Sullivan *et al.*, 2023, unpublished results) are shown in the first and second panels, respectively (top to bottom). The exons predicted by Vanacova *et al.* (2005), Wang *et al.* (2018), and the first genome annotation (2007) are shown in the following panels (3,4). The primers used for their validation by Wang *et al* (2018) are shown in the bottom panels (5,6).

**S3 Fig.**
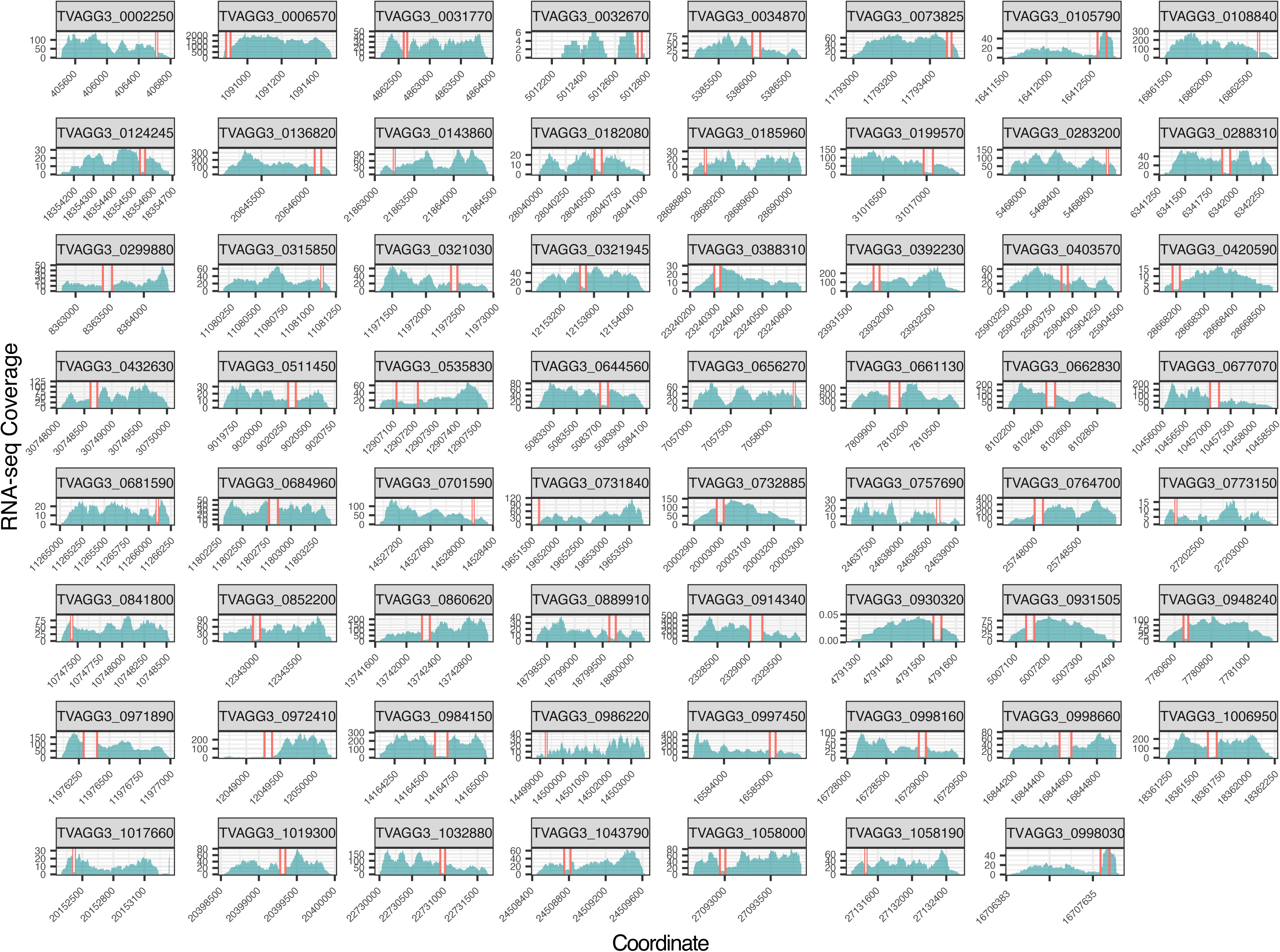
RNAseq coverage for the complete set of intron-containing transcripts in *T. vaginalis* G3. The intron sequences are delimited by vertical red lines, the RNAseq coverage is shown in cyan (y-axis), and the transcript length is indicated in the x-axis.

**S4 Fig.**
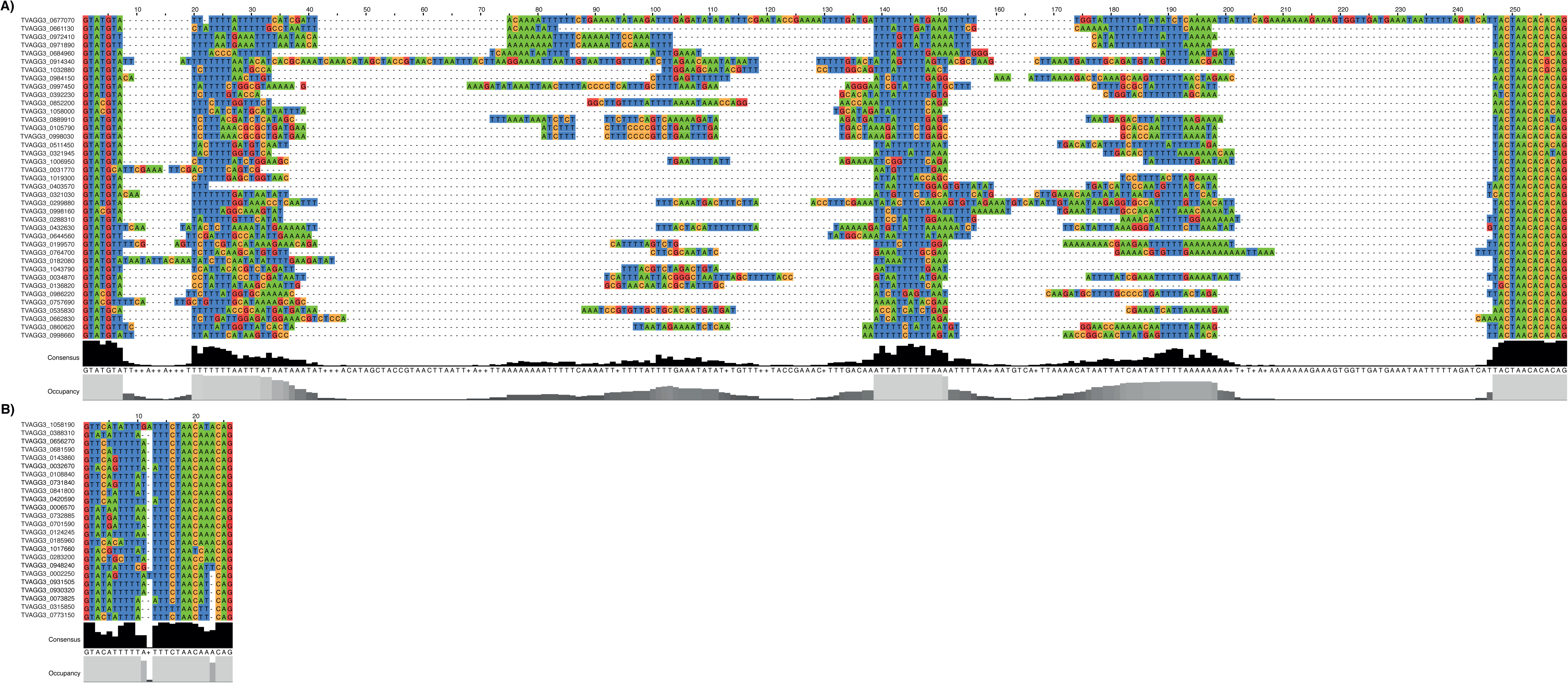
Multiple sequence alignment (MSA) of the 63 spliceosomal introns in *T. vaginalis*. The upper panel shows the MSA for the type A introns (length from 56-196 nucleotides), and the lower panel shows type B introns.

**S1 Table. The complete list of primers used for the PCR validation of the *T. vaginalis* and *T. stableri* introns.** Forward (top) and reverse (bottom) sequences and their main features are specified for all the primers. The length of the PCR product is specified in columns 6-7, and the length of the flanking intron is specified in column 8.

**S2 Table. Complete set of intron-containing genes in *T. vaginalis* and *T. stableri*.** All the data for both sets of spliceosomal introns, including their structural and functional annotation and the new corresponding gene IDs.

**S3 Table. The list of additional putative introns in *T. vaginalis* and *T. stableri* genomes.** These additional intron sequences were not validated by RNAseq evidence.

